# Estimating Body Segment Properties for Adults Across Diverse Body Morphologies: A Data-Driven Geometric Framework

**DOI:** 10.64898/2026.07.03.736346

**Authors:** Onorato d’Angelis, Chi Whan Choi, Hariharan Sureshkumar, Mario Merone, Simone V. Gill, Seungmoon Song

## Abstract

Accurate estimation of body segment inertial properties is essential for biomechanical analyses, yet commonly used scaling methods rely on limited datasets and do not generalize well across diverse adult body morphologies. We developed a data-driven framework that estimates segment lengths, masses, centers of mass, and moments of inertia using regression models trained on large anthropometric datasets (ANSUR II and NHANES) combined with a geometric representation of 16 body segments. The framework uses height, weight, and sex as primary inputs and incorporates waist and hip circumferences or other length and cross-sectional measurements when available to refine body-shape predictions. For individuals with obesity, additional geometric rules redistribute excess mass based on segment-specific volume changes. The resulting models reproduced segment lengths, cross-sectional dimensions, and lumped segment masses within the ranges observed in the training datasets and outperformed published regression equations, particularly at higher body mass index (BMI) values. To promote broad adoption, we provide an open-source API in Python that performs the full parameter estimation using the trained models. This framework offers an accurate and accessible method for estimating adult body segment properties across a wide range of body sizes and shapes, supporting improved motion analysis, musculoskeletal simulation, and clinical biomechanics.

## 1 Introduction

Accurately estimating the inertial properties of body segments is critical for biomechanical motion analysis [1], which are essential for movement rehabilitation [2, 3], sports science [4], and assistive devices [5, 6]. The dimensions, mass, center of mass (COM), and moments of inertia of individual body segments are required in analyzing the dynamics of motions. In typical motion analyses, such as gait analysis, motion capture data and external force measurements (e.g., ground reaction forces from force plates) is used to estimate the kinematics and dynamics of the recorded movement. The estimated kinematics and dynamics are often used in muscle dynamics analyses [7, 8], further informing biomechanical models for clinical and engineering applications. Several steps in these processes, such as inverse dynamics, muscle tuning, and analyzing muscle dynamics, depend on accurate estimates of body segment inertial properties. However, these estimations are often scaled from nominal values derived from anthropometric data collected decades ago from a small group of healthy individuals with average body types [9, 10]. Moreover, there is no straightforward method to validate the accuracy of these segment inertia estimations using motion data [11], making it challenging to ensure their reliability in diverse populations.

Directly measuring or accurately estimating body segment inertial properties for individuals is inherently challenging. Segment property data in the literature [12, 13] typically comes from two approaches: cadaver studies and volume-based estimation. Cadaver studies directly measure full body segment properties (i.e., dimensions and volume, COM, and moments of inertia) from cadaver dissections [10, 14]. While providing direct measures, this approach is not only infeasible for motion study participants, but also involves limitations arising from potential differences between living and deceased tissues. Volume-based approaches generally require specialized equipment, relying on measurements of segment volumes using techniques such as X-ray [15], dual-energy X-ray absorptiometry (DEXA) [16, 17], 3D laser scanning [18, 19], 3D photogrammetry [20], CT [21], and MRI [22]. The inertial properties are calculated based on the measured volume and presumed densities of different body components. Due to the limitations or challenges of these techniques, the studies often report segment properties of certain body parts, instead of all the body parts, and are conducted for people usually less than 100 with homogeneous ethnic backgrounds. The challenges of these approaches limit their use in common motion analysis studies to directly estimate segment properties of individuals participating in motion studies.

In practice, the difficulty of direct measurement has led researchers to estimate body segment properties using simplified geometric or regression models derived from limited data. Geometric models represent body segments as simple shapes, such as cylinders, cones, and ellipsoids from which the inertial properties are calculated [23,24]. Regression models represent the relationships between easily measurable subject anthropometric data, such as height and weight, and segment properties derived from cadaver studies or volume-based estimations [9, 10, 25, 26]. While these regression models provide an easy means of estimating segment properties of arbitrary human subjects, they are limited by the quality and quantity of the dataset from which they were derived, and there is no strong consensus on which model is the most reliable (as further analyzed in Section 2). As mentioned, most datasets that report comprehensive segment properties are often limited to small groups, ethnically homogeneous groups, and individuals with typical body types, excluding obese populations. Widely used motion analysis software, such as Visual3D [27], OpenSim [8], and AnyBody [28], offer customization options when additional data are available. However, by default, for most common motion analysis applications, these tools scale segment properties using simplified methods based on subject height and weight, with further scaling based on segment length from motion capture data. For instance, in Visual3D [27], segment lengths are derived from motion capture data, and segment masses are scaled from the default proportion to total body mass based on a cadaver study [10], published in 1955, based on data from eight white males, mostly under 60 kg. Moments of inertia are then calculated using geometric models of body segments proposed in 1964 [23].

There is a significant opportunity to utilize large anthropometric databases encompassing diverse body types and racial backgrounds to improve the estimation of body segment properties across different heights and weights for both sexes. For instance, AddBiomechanics [29], an open-source motion analysis tool, uses the ANSUR II dataset [30], which includes segment lengths from over 6,000 U.S. military adults, as a statistical prior for scaling segment lengths based on motion capture data. While, to our knowledge, there is no publicly available large database that includes comprehensive segment properties such as mass and moments of inertia of individual segments, several large datasets report segment lengths [30], segment circumferences [30,31], and estimated mass of certain body segments [32]. These datasets represent thousands of individuals across different heights, weights, sexes, and racial backgrounds. Thus, there is an opportunity to utilize these datasets to estimate segment properties, including full inertial properties of individual segments, for more accurate and inclusive biomechanical analyses.

Here, we propose a novel approach to estimate comprehensive properties of individual body segments, including those of obese populations, by incorporating large anthropometric datasets. Our framework uses machine learning regression models trained with large datasets, which take the height, weight, and sex of the target subject as inputs, to estimate segment lengths, certain circumferences, and the masses of major body segments. These estimated properties are processed (e.g., by using regression models to derive smaller body segment properties and by incorporating optional anthropometric inputs), and then used to determine geometric models of individual segments, which are subsequently used to calculate the COM and moments of inertia. The main contributions of this project are as follows:

- An improved segment scaling method that integrates large, inclusive datasets and geometric modeling, applicable to both typical and obese populations.
- A demonstration that widely used regression models of segment properties are often inaccurate when compared to available large datasets.
- Confirmations of key anthropometric relationships, including that segment lengths depend only on height and sex rather than body weight, and that waist and hip circumferences well capture major body-shape variation.
- An open-source Python API made available for developers and researchers to integrate our methods into their workflows.

## 2 Datasets and prior regression models for segment properties

In this section, we (i) summarize the large, publicly available datasets used in this study, (ii) review commonly used regression models for estimating body-segment properties, and (iii) report their estimation accuracy against those datasets. Full details of the evaluation protocol, metrics, and stratified results are provided in Supplementary A. We also highlight the best performing model, which we incorporate into our framework and use as the primary benchmark.

We used two large, publicly available datasets that report anthropometric measurements and estimate relevant segment properties: ANSUR II [30] and NHANES [32]. Both datasets include height, weight, and sex, along with a range of additional anthropometric variables. ANSUR II (the Anthropometric Survey of U.S. Army Personnel) provides detailed body measurements from 4,082 males and 1,986 females, from which we extracted segment lengths and cross-sectional dimensions. NHANES (National Health and Nutrition Examination Survey) includes body composition data obtained via dual-energy X-ray absorptiometry (DEXA) for 5,288 males and 5,055 females; from NHANES we used mass estimates for the torso, head/neck, legs, and arms. We did consider incorporating circumference data from the AWS BodyM dataset [31], but ultimately relied on ANSUR II, as we could not find precise definitions of the measurement sites in BodyM and ANSUR II provides broader coverage across the BMI spectrum.

Table 1 summarizes the predictive accuracy of published regression models when evaluated against ANSUR II and NHANES. We surveyed models that estimate lengths, masses, centers of mass (COM), and moments of inertia from basic anthropometry (e.g., height and body mass) [9, 25, 26, 33]. Across most strata (segment, sex, and racial/ethnic groups), the Shan–Bohn German model demonstrated the lowest or near-lowest errors. We therefore designate it as our *literature-best (SB–G) baseline* for benchmarking, and we incorporate its mass-partitioning within our framework (Section 3.2).

**Table 1:** Summary of method-average mean absolute errors (percentage) for segment length and segment mass. Values correspond to the method-average mean absolute errors (MAE, %) reported in Supplementary Tables S2–S4.

| Method | Segment length MAE (%) |  | Segment mass MAE (%) |  |
| --- | --- | --- | --- | --- |
|  | Male | Female | Male | Female |
| Contini | 6.88 | 5.09 | – | – |
| Shan Asian | 6.35 | 5.41 | 13.75 | 15.07 |
| Shan German | 4.94 | 4.58 | 16.24 | 13.09 |
| Finch | – | 10.77 | – | – |
| Zatsiorsky | – | – | 67.28 | 70.37 |

## 3 Methods

Our framework provides estimations of segment properties for male and female adults across a wide range of heights and weights (BMI: 15–40 kg/m^2^ for both males and females; males: 1.50–2.00 m, and females: 1.40–1.80 m) by incorporating four methodological components: a geometric model of the entire body, data-driven anthropometric parameter estimation, obese-specific adjustments, and final rescaling (Fig. 1). Given the target subject’s height, weight, and sex, the baseline process includes: (1) estimating segment lengths, masses, and cross-sectional properties using data-based models (e.g., data-driven estimators trained on large datasets), and (2) determining the geometric model with these parameters to calculate the center of mass and moments of inertia. For obese cases (BMI ≥30 kg/m^2^), we represent each segment with a core-plus-extension geometry, distributing the excess body mass according to the volume of the extended segments. Additionally, if optional inputs such as segment lengths and cross-sections are provided, the geometric model is further refined to incorporate these inputs.

**Figure 1:**
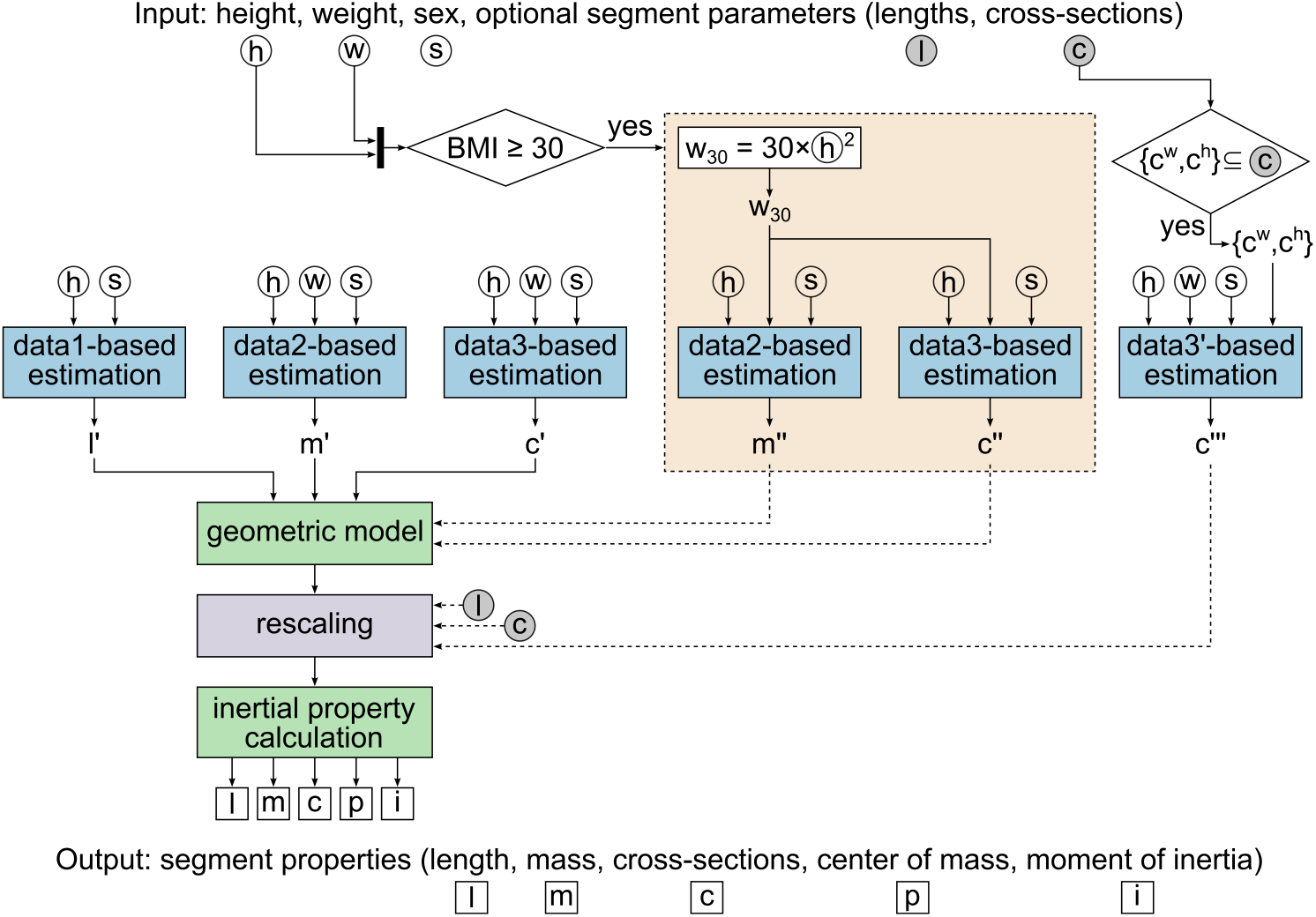
Overview of the proposed framework for estimating body segment properties.

Each of these methodological components involves specific intricacies that are described in the following subsections. First, the geometric model is introduced, as all parameters are defined within it (3.1). Next, we outline the data-based estimations that set the lengths, masses, and cross-sections of the geometric model (3.2). We then detail the obese-specific adjustments (3.3) and conclude with the final rescaling procedure (3.4).

### 3.1 Geometric model

The geometric model is central to our framework, combining all anthropometric parameters to fully define the body segments and calculate their inertia properties. Our geometric model consists of 16 segments, including the head/neck, upper torso, middle torso, lower torso, thighs, shanks, feet, upper arms, forearms, and hands (Fig. 2). Each segment is represented using either an ellipsoid, a frustum of an elliptical cone, an elliptical cone, or a combination of these shapes. In total, the model includes 41 parameters: 13 lengths, 10 masses, and 18 cross-sectional properties. Once these parameters are estimated primarily from the data-based models or using novel estimation techniques (see Subsection 3.2), the COM and moments of inertia are calculated, assuming uniform density.

**Figure 2:**
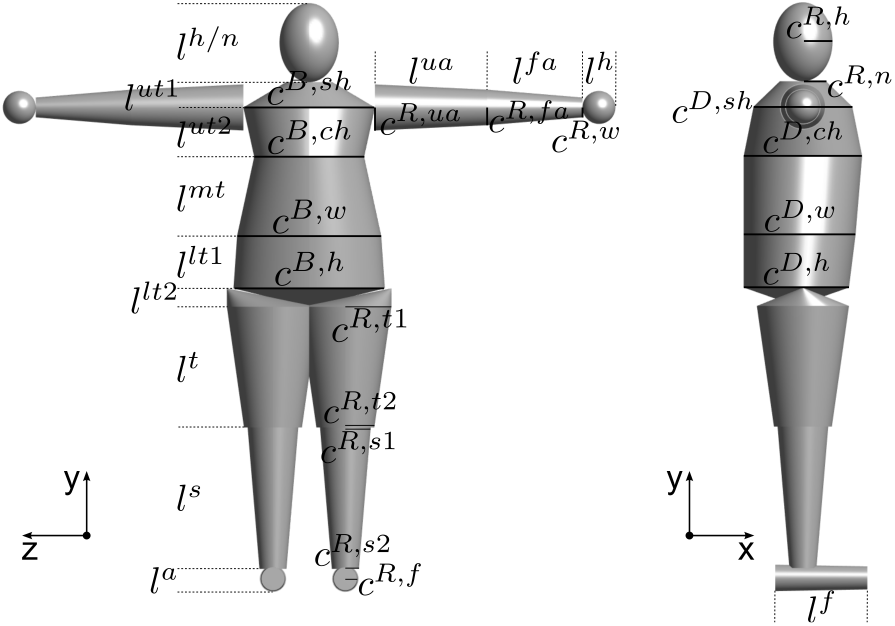
Geometric model and parameters. The 3D model illustrates all body segments and the dimensional parameters used to calculate segment centers of mass and moments of inertia. In addition to the shown length and cross-section parameters, each segment is also defined by its mass (*m*^*h/n*^, *m*^*ut*^, *m*^*mt*^, *m*^*lt*^, *m*^*t*^, *m*^*s*^, *m*^*f*^ , *m*^*ua*^, *m*^*fa*^, *m*^*h*^), which are not displayed.

**Figure 3:**
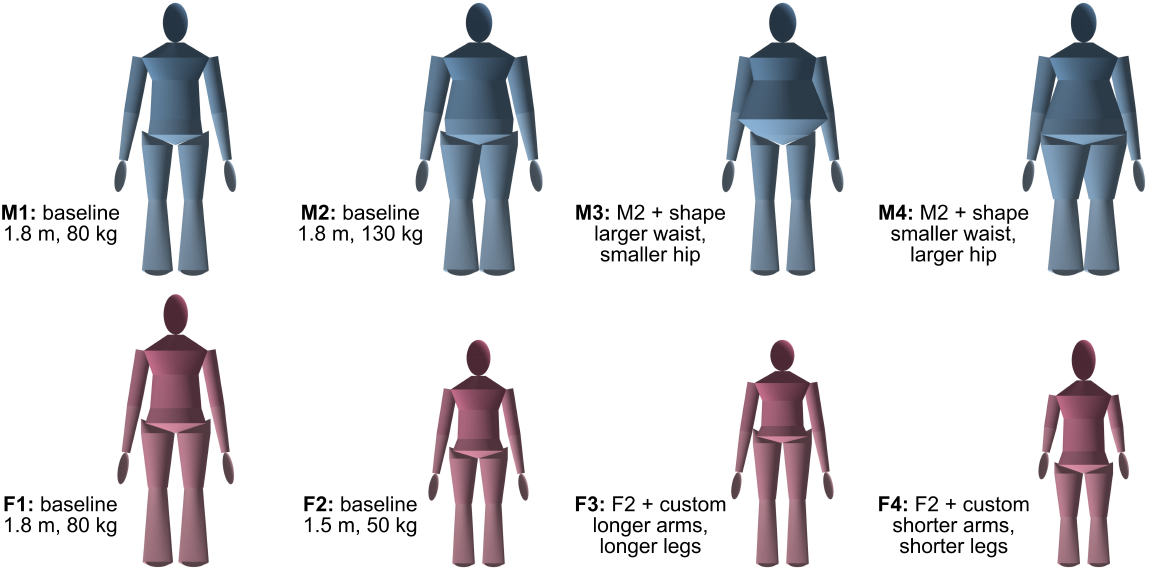
Example geometric body models generated by the proposed API. The models illustrate the range of individualized geometries that can be created using the API modes shown in Fig. 1. The first row (M1–M4) shows male models, and the second row (F1–F4) shows female models. In the baseline mode, segment lengths, masses, and cross-sectional parameters are estimated from height, body mass, and sex. In the shape mode, waist and hip circumferences are additionally provided to refine body-shape estimates. In the custom mode, user-specified segment lengths and/or cross-sectional measurements are incorporated into the model. M1 shows a baseline male model (1.8 m, 80 kg), and M2 shows a baseline male model with the same height but greater body mass (1.8 m, 130 kg). M3 and M4 are generated with the same height, body mass, and sex as M2 but using the shape mode, with M3 specifying a larger waist circumference and smaller hip circumference and M4 specifying a smaller waist circumference and larger hip circumference. F1 shows a baseline female model with the same height and body mass as M1; while the visual differences are subtle, sex-specific estimation yields slightly different body geometries. F2 shows a shorter and lighter baseline female model (1.5 m, 50 kg). F3 and F4 are generated from F2 using the custom mode, with longer arms and legs specified for F3 and shorter arms and legs specified for F4.

Our geometric model includes more parameters than conventional models to better represent diverse body types. Previous geometric models [23, 34, 35] were typically parameterized for straightforward derivation from a limited set of anthropometric variables reported in public datasets. However, these models are often insufficient for capturing the geometric characteristics of individuals with diverse or obese body types. In addition to incorporating 17 cross-sectional parameters available in the ANSUR II dataset, we introduced the shoulder depth (*c*^*D,sh*^) parameter to capture the anterior-posterior mass distribution trend reported for individuals with obesity [36] (see also Section 3.3).

A detailed description of the geometric model, including the rationale for each assumption and the equations for calculating the COM and moments of inertia, is provided in Supplementary B.

### 3.2 Data-based parameter estimation

The segment length, mass, and cross-section parameters that define the geometric model are estimated primarily using the ANSUR II [30] and NHANES [32] datasets. For the majority of segment parameters, these datasets provided sufficient coverage across the target BMI range, and we trained regression models to estimate them. For each parameter, we evaluated five regression approaches (linear regression, support vector regression, Lasso regression, decision trees, and random forests) and selected the method with the lowest root mean square error (RMSE) in 5–fold cross-validation. Full model specifications and performance results are reported in Supplementary C. For parameters without sufficient data, we used novel estimation techniques informed by existing regression models from smaller datasets or heuristics derived from prior studies.

All segment lengths are estimated using regression models trained on the ANSUR II dataset (Fig. 1, data1-based estimation). These models use height and sex as inputs. We confirmed, by comparing models with and without weight as an additional input, that including weight did not yield a statistically significant improvement (Supplementary D), consistent with previous reports [9].

To estimate segment masses, we combined regression models trained on the NHANES dataset with established models from the literature (Fig. 1, data2-based estimation). The models use height, weight and sex as inputs. First, the NHANES-based regression models provide estimates of lumped masses for the trunk, legs, and arms. These lumped masses are then distributed to individual segments using the SB–G regression model (our literature-best reference model; see Section 2).

Cross-sectional parameters are primarily estimated using regression models trained on the ANSUR II dataset (Fig. 1, data3-based estimation). The models use height, weight, and sex as inputs for the primary estimations, while additional models incorporating waist and hip circumferences are trained for cases where those measures are available, enabling refined rescaling of the segments (data3^*′*^-based estimation; Section 3.4). Training these additional models with waist and hip circumferences was motivated by two observations: 1) feature-selection analyses showed that these two measures account for the majority of cross-sectional body-shape variation (Supplementary D), and 2) their inclusion aligns with established practice that uses waist and hip measurements as primary descriptors of body type, such as apple- and pear-shaped profiles [37]. All cross-sectional parameters are estimated from the ANSUR II-based regression models, except for shoulder depth (*c*^*D,sh*^), which is set equal to chest depth (*c*^*D,ch*^) for BMI*<*30 kg/m^2^ and remains constant for higher BMI values (see Section 3.3 for rationale).

### 3.3 Obese-specific adjustments

For obese cases (BMI ≥30 kg/m^2^), the framework employs a two-stage approach to better represent the additional body mass associated with adiposity. BMI=30 kg/m^2^ is chosen as the reference for constructing the base geometric model, assuming that any mass beyond this threshold primarily represents fat, which differs in density from the core segments. First, the framework estimates the mass and cross-section parameters for all segments as if the individual had the same sex and height but BMI=30 kg/m^2^ (base segments; boxed in orange in Fig. 1). It then re-estimates the segment cross-sections for the target BMI (≥30 kg/m^2^) to determine the increased segment volumes, or extended segments. Using the NHANES-based regression models within the data2-based estimation, the framework computes the surplus masses of the lumped trunk, leg, and arm segments, which are subsequently distributed to individual segments in proportion to their relative increases in volume.

Additional geometric rules were encoded in the model and its parameterization to better capture the characteristic body shape of individuals with obesity [36]. First, the posterior boundaries of the middle and lower torso are aligned with that of the chest (whose depth is determined from the ANSUR II-based regression model) so that the torso geometry is skewed anteriorly, reflecting the forward mass distribution typical of obesity. Second, the shoulder depth (*c*^*D,sh*^), which is not available in the dataset, is assumed to follow the chest depth (*c*^*D,ch*^) up to a BMI of 30 kg/m^2^ and to remain fixed for higher BMI values. This constraint reflects the typical pattern in obesity, where additional body mass beyond this threshold predominantly accumulates in the lower torso rather than the upper body.

### 3.4 Optional inputs, rescaling, and calculation of segment

Users may optionally provide additional anthropometric measurements, including selected segment lengths, circumferences, breadths, depths, and length/height measures. These optional inputs are incorporated directly when they correspond to internal geometric parameters, or indirectly when they are used to estimate derived model dimensions. When waist and hip circumferences (*c*^*w*^, *c*^*h*^) are provided, all other cross-sectional parameters are re-estimated through the data3^*′*^-based estimation to maintain geometric consistency, unless any of them are explicitly specified by the user.

Rescaling is conducted after incorporating all optional inputs. Even when no additional inputs are provided, this step is necessary to ensure that the total body height and mass match the input values, since individual segment parameters are independently estimated from regression models and would not sum precisely to the input height and weight. Segment lengths are rescaled by the ratio 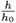, where *h* is the input height and *h*_0_ is the current height, so that their total matches the input value. Segment masses are then updated to preserve each segment’s density relative to the whole-body density as

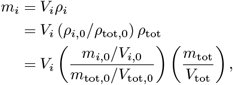

where *i* indexes individual body segments; *m*_*i*_, *V*_*i*_, and *ρ*_*i*_ are the updated mass, volume, and density of segment *i*, respectively; and the subscript 0 denotes the corresponding values before rescaling. *m*_tot_ is the input body mass (*w*), *V*_tot_ is the updated total body volume, and *ρ*_tot_ = *m*_tot_*/V*_tot_ is the updated whole-body density. Similarly, *m*_tot,0_, *V*_tot,0_, and *ρ*_tot,0_ = *m*_tot,0_*/V*_tot,0_ denote the total body mass, volume, and density before rescaling.

Once all segment lengths, masses, and cross-sectional values are updated, the COM and moments of inertia are calculated from the geometric model, which together constitute the complete set of segment parameters provided by our framework.

## 4 Results

Our framework was successfully implemented as designed, integrating data-driven parameter estimation with a geometric model to generate full segment inertial properties across adult BMI ranges from 15 to 40 kg/m^2^ in both males and females. The approach accurately reproduces segment lengths, masses, and cross-sectional parameters within the ranges represented in ANSUR II and NHANES, and it scales smoothly with changes in height, weight, and optional anthropometric inputs. The full implementation is available through Python API, and the source code is provided on GitHub (https://github.com/neumovelab/BodySegmentEstimator) [38].

The data-based estimations are well trained, as the estimated segment lengths, masses, and cross-sectional parameters closely matched the corresponding values reported in the ANSUR II and NHANES datasets (full RMSE results are provided in Supplementary C). Figure 4 illustrates selected parameters and their aggregated values, showing strong agreement between our model estimations and the large-scale datasets (an expected outcome since the models were trained using these data). In contrast, the literature-best (SB–G) baseline [26] deviates substantially from the large datasets (RMSE results in Supplementary A), reflecting biases introduced by its limited training population of only 25 subjects. For example, the SB–G model exhibits notable variance in head (male), torso, and leg (female) lengths, indicating high dependencies on body weight, whereas our analysis indicates that segment lengths are not statistically affected by weight (at least within these datasets) and therefore height and sex alone are used as inputs. Also, the lumped segment masses predicted by the SB–G model diverge markedly, especially at higher BMI values.

**Figure 4:**
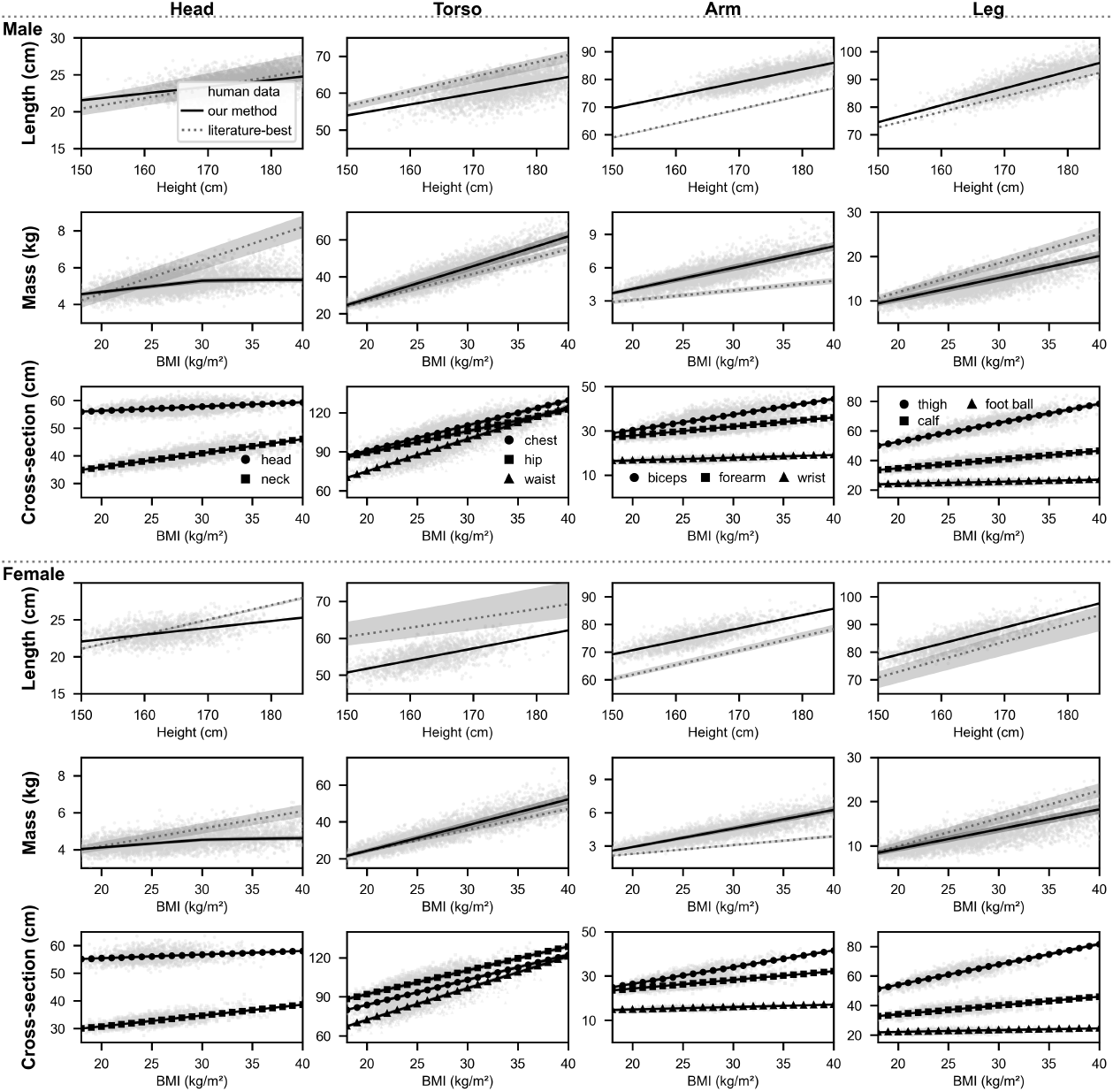
Estimation of segment length, mass, and cross-sectional parameters. Predictions from our framework (solid lines), the literature-best SB–G regression model [26] (dotted lines), and human data from the ANSUR II and NHANES datasets (gray markers) are shown for males (top) and females (bottom). Shaded regions indicate the BMI range used for length estimation (15–43 kg/m^2^) and the height ranges used for mass and cross-sectional estimation (males: 136– 199 cm; females: 135–186 cm). An ideal fit follows the central trend of the data cloud.

The masses of the torso segments are reasonably distributed across the BMI range with the obese-specific adjustments incorporated in our framework. Figure 5 compares the predicted masses of the torso segments from our framework with those from a version without obese-specific adjustments (i.e., without using the base and extended segments; boxed in orange in Fig. 1) and with the literature-best (SB–G) baseline [26]. As intended, our framework allocates proportionally more mass to the lower torso for BMI ≥30 kg/m^2^, consistent with the characteristic mass distribution observed in individuals with obesity [36].

**Figure 5:**
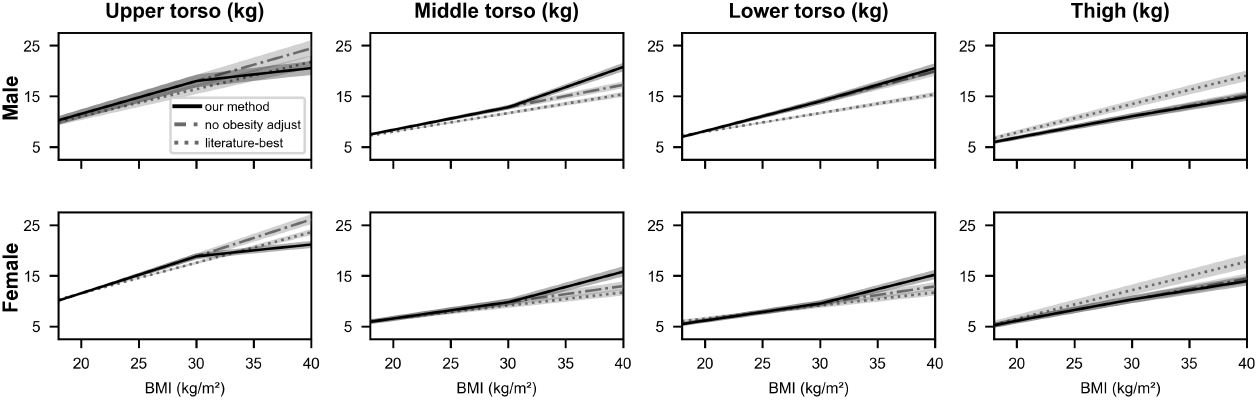
Torso and thigh segment masses. The figure compares estimates from our framework (solid lines), our framework without obese-specific adjustments using dual base and extended segments (dash–dot lines), and the literature-best SB–G regression model [26] (dotted lines). Our framework reproduces the expected redistribution of mass toward the middle and lower torso with increasing BMI.

Direct validation of individual segment masses and inertial properties remains limited, particularly for individuals with obesity, as few studies report comprehensive segment-level data. To provide a representative comparison, we evaluated our framework against a version without obesity-specific adjustments and the literature-best baseline [26] using segment inertial parameters from an individual with central adiposity estimated via dual-energy X-ray absorptiometry (DEXA) and 3D geometric reconstruction [36]. Our framework yielded the lowest error (RMSE = 2.46 kg, *R*^2^ = 0.98), outperforming both the version without obeity adjustments (RMSE = 2.77 kg, *R*^2^ = 0.97) and the literature-best baseline (RMSE = 5.22 kg, *R*^2^ = 0.91). Moreover, our framework reproduced the characteristic anterior mass distribution and greater lower-torso segment masses observed in that dataset, whereas the baseline substantially underestimated these effects.

Figure 6 illustrates the effect of incorporating waist and hip circumferences as optional inputs to characterize different obese body types. The baseline represents the default model without optional inputs, whereas the “more apple-like” and “more pear-like” cases correspond to relatively larger waist or hip circumferences, respectively, compared with the baseline. As expected, increasing the waist circumference (more apple-like type) results in greater mass allocation to the upper and middle torso segments, whereas increasing the hip circumference (pear-like type) increases the mass of the thighs. These trends are consistent across sexes and align with reported distinctions in fat distribution patterns between apple- and pear-type obesity.

**Figure 6:**
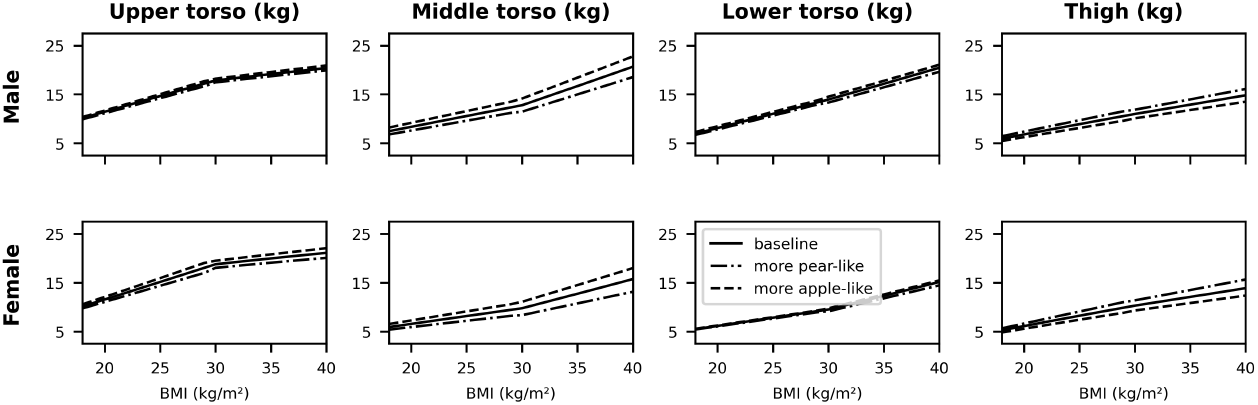
Torso and thigh segment masses with optional waist and hip circumference inputs. The figure shows estimates obtained using only the essential inputs (height, weight, and sex; solid lines) and those obtained with additional waist and hip circumferences. The optional circumferences were varied to represent a more pear-like body type (dash–dot lines) and a more apple-like body type (dashed lines). More apple-like inputs increase upper and middle torso mass, whereas more pear-like inputs increase thigh mass.

## 5 Discussion

We developed a data-driven framework that estimates comprehensive body segment properties across adult BMI ranges (15–40 kg/m^2^) in both males and females. By combining regression models trained on large, diverse datasets (ANSUR II and NHANES) with a geometric model that adapts to individual morphology, the framework provides accurate segment masses, centers of mass, and moments of inertia from minimal inputs (height, weight, and sex) with optional anthropometric refinements. Unlike conventional regression or proportional scaling methods, which extrapolate from limited cadaver data and underrepresent obese individuals, our approach generalizes across a broad BMI range and better captures characteristic mass distributions. Despite its greater complexity relative to prior methods, the framework was designed to remain interpretable by incorporating explicit geometric representations and interpretable regression models. These improvements enable more realistic modeling of human body dynamics, particularly for populations not adequately represented in existing models. This publication is accompanied by open Python API, facilitating broad accessibility for research and clinical applications.

In developing the framework, we conducted analyses that formally confirmed several anthropometric relationships not clearly established in prior literature (Supplementary A and D). First, segment lengths were not statistically influenced by body weight (despite its inclusion in previous regression models [9, 26]) indicating that height and sex alone are sufficient for estimating segment lengths in adults. Second, waist and hip circumferences emerged as key predictors of body-shape variation, effectively representing broader cross-sectional characteristics through feature selection. This finding supports their conventional use as descriptors of body type (e.g., apple- and pear-shaped) [39]. Finally, all literature regression models we evaluated [9, 25, 26, 33], including those widely used in biomechanics, deviated markedly from large population datasets, showing systematic biases across both obese and normal-weight ranges. These results highlight the need to re-evaluate legacy scaling models that remain embedded in common motion analysis software.

While the proposed framework represents a substantial improvement over previous models, several limitations should be acknowledged, primarily reflecting the lack of comprehensive, standardized datasets for human body segment properties. First, the ANSUR II dataset [30] comprises U.S. military personnel and, while demographically diverse in race and BMI, may not fully represent the characteristics of the general population, which may introduce representation bias. We considered incorporating additional datasets such as BodyM [31] to mitigate this bias; however, the available documentation does not clearly define the anatomical measurement sites for the reported cross-sectional variables, limiting its compatibility with our framework. Second, the NHANES dataset provides estimated masses for the trunk, arms, and legs but lacks detailed segment-level data, requiring the use of regression models from the literature [26] trained on smaller datasets to distribute mass among individual segments. Third, current datasets contain few samples with higher BMI values, particularly above 40 kg/m^2^, restricting validations for severely obese individuals. These limitations could be addressed as more extensive and standardized datasets become available, enabling a more accurate and streamlined estimation framework. With future datasets, additional features such as age could also be incorporated as key predictors, as age likely influences body composition and mass distribution across segments.

Beyond leveraging improved datasets as they become available, future work could proceed along two main directions. First, more advanced geometric representations could be incorporated into the framework by integrating comprehensive body-shape measurement techniques. Full-body mesh models such as SMPL [40], widely used in computer vision for reconstructing 3D human shape and pose from images or videos [41], could enable personalized calibration of body geometry and replace our current estimates of selected cross-sectional parameters (Fig. 1, data3-based estimation). Second, future studies should apply the framework within motion analysis and predictive neuromechanical simulations [42] to assess its efficacy relative to existing scaling approaches for estimating segmental dynamics, joint loading, and muscle forces, and for simulating overall movement. Integration into widely used tools such as Visual3D [27], OpenSim [8], AnyBody [28], and MyoSuite [43, 44] would facilitate such validation and promote broader adoption within the biomechanics community.

