## Supplementary Material for "Estimating Body Segment Properties for Adults Across Diverse Body Morphologies: A Data-Driven Geometric Framework"

**Contents**


- Supplementary A: Datasets and prior regression models
- Supplementary B: Geometric model
- Supplementary C: Regression models
- Supplementary D: Extended results


### Supplementary A. Datasets and prior regression models

This section provides supplementary details for Section 2 of the main article. It describes the datasets used to evaluate prior regression models for body segment parameters and reports the stratified error metrics underlying the summary presented in Table 1 of the main text.

#### A.1 Datasets

The segment length, mass, and cross-section parameters that define the geometric model are estimated primarily using large public anthropometric datasets. Parameters with sufficient data available across the target BMI range are estimated using regression models trained on these datasets. For parameters that are not directly available, we employed estimation techniques based on existing regression models from smaller datasets or heuristics derived from prior studies.

The ANSUR II dataset [1] is a comprehensive collection of anthropometric measurements from over 6,000 adult U.S. military personnel, consisting of 4,082 men and 1,986 women. It includes 93 measurements and provides a detailed overview of body size and shape for a diverse population aged 17 and older. The dataset includes several ethnic groups such as White, Black, Hispanic, Asian, and Native American. We used ANSUR II to build regression models for body segment lengths and cross-sectional parameters (e.g., breadth, depth, and circumferences), and to validate the segment-length components of previously published regression equations.

The NHANES dataset [2] provides dual-energy X-ray absorptiometry (DXA)–based estimates of body segment masses for the trunk, arms, legs, and head. In this study, we used these DXA-derived segment masses to train regression models that predict lumped segment masses from height, weight, and sex. The NHANES samples used for these analyses include 5,288 males and 5,055 females, spanning a wide range of BMI values, and were drawn from the 2011–2018 survey cycles. These data enabled both the evaluation of previously published mass-regression equations and the development of the data-driven mass estimators incorporated into our framework.

Table S1 summarizes the BMI distributions for ANSUR II and NHANES by sex.

**Table S1:** BMI classification frequencies for ANSUR II and NHANES, stratified by sex.

| Dataset | Sex | Underweight | Normal | Overweight | Obesity I | Obesity II | Obesity III |
|---|---|---|---|---|---|---|---|
| ANSUR II | Male | 17 (0.4) | 1053 (25.8) | 1904 (46.6) | 921 (22.6) | 169 (4.1) | 18 (0.5) |
| | Female | 20 (1.0) | 929 (46.8) | 847 (42.6) | 171 (8.6) | 17 (0.9) | 2 (0.1) |
| NHANES | Male | 107 (2.0) | 1595 (30.2) | 1949 (36.9) | 1039 (19.6) | 384 (7.3) | 214 (4.0) |
| | Female | 161 (3.2) | 1628 (32.2) | 1361 (26.9) | 968 (19.1) | 537 (10.6) | 400 (8.0) |

*Values are reported as count (percent).*
*BMI category thresholds:* Underweight ($< 18.8$ kg/m$^2$), Normal ($< 25$), Overweight ($< 30$), *Obesity I* ($< 35$), *Obesity II* ($< 40$), *Obesity III* ($\geq 40$).

#### A.2 Evaluation of prior regression models

In the existing literature, numerous regression equations have been developed to derive body segment parameters (BSPs) from height and weight. To determine which equations to adopt as a reference in this study, we considered models that (i) require only stature and body mass as inputs, (ii) provide parameters for both segment length and mass, and (iii) include separate equations for males and females. Under these criteria, we evaluated the methods of Contini [3], Shan and Bohn (German and Asian equations) [4], Finch (female-specific) [5], and Zatsiorsky [6].

To evaluate the performance of existing regression equations for predicting body segment parameters, we analyzed segment lengths using the ANSUR II dataset and lumped segment masses using NHANES. For the length evaluation, subjects labeled as *Unknown* ethnicity were excluded, and the Interquartile Range (IQR) method was applied separately within each ethnic group to identify and remove outliers. Each regression method was tested independently across ethnic groups and across multiple segments of the arms, legs, and torso. We computed the mean absolute error (MAE) for each segment and summarized percentage errors for five representative segments corresponding to the measurement points used in the main paper: foot, hand, forearm, shank, and head. Tables S2 and S3 report the resulting percentage errors for males and females, respectively. For the mass evaluation, lumped segment masses (arms, legs, trunk, and head) were assessed using NHANES, and the percentage errors for each method are summarized in Table S4.

**Table S2:** Mean absolute error (percentage) in segment length by ethnicity, method, and body segment (male).

| **Ethnicity** | **Contini Method (%)** | | | | | **Shan Asian (%)** | | | | | **Shan German (%)** | | | | |
|---|---|---|---|---|---|---|---|---|---|---|---|---|---|---|---|
| | Foot | Hand | Forearm | Shank | Head | Foot | Hand | Forearm | Shank | Head | Foot | Hand | Forearm | Shank | Head |
| White | 2.58 | 17.37 | 4.02 | 8.29 | 4.10 | 5.56 | 9.33 | 2.89 | 3.72 | 10.74 | 4.40 | 3.47 | 3.00 | 7.59 | 5.55 |
| Black | 4.00 | 12.76 | 5.64 | 4.87 | 4.66 | 8.25 | 7.19 | 6.82 | 2.81 | 13.20 | 6.98 | 3.15 | 7.20 | 4.40 | 7.71 |
| Hispanic | 2.90 | 16.42 | 4.32 | 8.23 | 3.85 | 5.97 | 10.38 | 3.99 | 3.52 | 8.04 | 5.17 | 3.03 | 3.70 | 6.47 | 5.63 |
| Asian | 2.65 | 16.98 | 4.26 | 11.36 | 4.81 | 4.14 | 8.05 | 3.63 | 6.83 | 5.45 | 3.75 | 3.42 | 3.27 | 8.46 | 4.45 |
| Native American | 3.44 | 17.47 | 4.64 | 8.46 | 3.22 | 7.26 | 11.98 | 3.18 | 3.21 | 11.93 | 6.00 | 4.49 | 3.78 | 8.17 | 5.98 |
| Pacific Islander | 3.01 | 14.33 | 4.28 | 9.46 | 3.32 | 6.67 | 6.66 | 4.57 | 4.85 | 6.27 | 5.96 | 2.27 | 3.80 | 7.30 | 4.54 |
| Unknown | 1.41 | 13.33 | 3.81 | 3.70 | 2.84 | 4.97 | 5.27 | 6.36 | 0.89 | 7.83 | 4.13 | 5.33 | 4.45 | 2.18 | 3.84 |
| **Average** | 2.86 | 15.52 | 4.42 | 7.76 | 3.82 | 6.12 | 8.41 | 4.49 | 3.69 | 9.06 | 5.20 | 3.59 | 4.17 | 6.37 | 5.38 |
| **Method Average** | 6.88 | | | | | 6.35 | | | | | 4.94 | | | | |

**Table S3:** Mean absolute error (percentage) in segment length by ethnicity, method, and body segment (female).

| **Ethnicity** | **Contini Method (%)** | | | | | **Shan Asian (%)** | | | | | **Shan German (%)** | | | | | **Finch (%)** | | | | |
|---|---|---|---|---|---|---|---|---|---|---|---|---|---|---|---|---|---|---|---|---|
| | Foot | Hand | Forearm | Shank | Head | Foot | Hand | Forearm | Shank | Head | Foot | Hand | Forearm | Shank | Head | Foot | Hand | Forearm | Shank | Head |
| White | 3.13 | 3.22 | 11.00 | 6.33 | 4.24 | 4.26 | 3.52 | 4.99 | 2.98 | 5.44 | 3.44 | 4.08 | 3.86 | 6.71 | 4.41 | 20.88 | 4.98 | 11.14 | 12.20 | 4.43 |
| Black | 2.86 | 3.99 | 6.62 | 3.50 | 4.38 | 6.08 | 6.69 | 10.25 | 5.56 | 6.32 | 5.63 | 4.27 | 4.30 | 3.52 | 5.62 | 23.51 | 8.92 | 6.71 | 8.20 | 4.53 |
| Hispanic | 2.44 | 3.01 | 9.18 | 5.70 | 4.17 | 4.02 | 4.75 | 6.73 | 3.52 | 6.04 | 3.81 | 3.55 | 3.37 | 5.58 | 3.82 | 22.16 | 6.29 | 9.30 | 11.36 | 4.39 |
| Asian | 3.14 | 3.01 | 10.96 | 8.17 | 5.33 | 2.95 | 4.80 | 5.42 | 3.57 | 6.14 | 2.79 | 3.55 | 4.69 | 8.57 | 4.74 | 20.76 | 5.88 | 11.10 | 13.97 | 5.65 |
| Native American | 3.05 | 2.50 | 7.87 | 5.38 | 4.32 | 5.47 | 4.58 | 7.08 | 3.58 | 5.65 | 4.79 | 3.33 | 5.41 | 5.09 | 4.85 | 22.97 | 7.10 | 7.98 | 10.01 | 4.35 |
| Pacific Islander | 3.13 | 3.47 | 8.62 | 4.95 | 5.12 | 4.84 | 5.96 | 8.18 | 4.15 | 8.76 | 4.79 | 4.61 | 4.38 | 5.31 | 4.56 | 22.23 | 7.55 | 8.77 | 10.56 | 5.29 |
| **Average** | 2.96 | 3.20 | 9.04 | 5.67 | 4.59 | 4.60 | 5.05 | 7.11 | 3.89 | 6.39 | 4.21 | 3.90 | 4.34 | 5.80 | 4.67 | 22.09 | 6.79 | 9.17 | 11.05 | 4.77 |
| **Method Average** | 5.09 | | | | | 5.41 | | | | | 4.58 | | | | | 10.77 | | | | |

**Table S4:** Mean absolute error (percentage) in segment mass by ethnicity, method, and body segment (male and female).

| **Sex / Ethnicity** | **Zatsiorsky (%)** | | | **Shan Asian (%)** | | | **Shan German (%)** | | |
|---|---|---|---|---|---|---|---|---|---|
| | Arm | Leg | Trunk | Arm | Leg | Trunk | Arm | Leg | Trunk |
| **Male** | | | | | | | | | |
| White | 118.50 | 77.20 | 9.73 | 14.76 | 16.80 | 5.85 | 25.88 | 18.66 | 8.17 |
| Black | 119.01 | 67.84 | 4.72 | 19.23 | 21.51 | 12.32 | 29.55 | 11.86 | 4.11 |
| Asian | 120.89 | 76.08 | 9.16 | 12.73 | 17.38 | 6.71 | 24.81 | 16.79 | 6.54 |
| Other | 119.95 | 75.48 | 8.78 | 13.14 | 17.54 | 7.00 | 25.16 | 16.76 | 6.54 |
| **Average (male)** | 119.59 | 74.15 | 8.10 | 14.97 | 18.31 | 7.97 | 26.35 | 16.02 | 6.34 |
| **Method Average** | 67.28 | | | 13.75 | | | 16.24 | | |
| **Female** | | | | | | | | | |
| White | 130.86 | 66.02 | 8.49 | 30.08 | 5.97 | 3.86 | 25.14 | 11.14 | 5.64 |
| Black | 129.84 | 60.97 | 5.48 | 31.96 | 5.97 | 5.51 | 26.12 | 7.96 | 5.26 |
| Asian | 136.71 | 74.93 | 9.94 | 30.97 | 12.30 | 5.31 | 21.69 | 11.14 | 4.94 |
| Other | 136.32 | 74.98 | 9.83 | 31.45 | 12.23 | 5.25 | 22.21 | 10.92 | 4.95 |
| **Average (female)** | 133.43 | 69.23 | 8.44 | 31.12 | 9.12 | 4.98 | 23.79 | 10.29 | 5.20 |
| **Method Average** | 70.37 | | | 15.07 | | | 13.09 | | |

Although several methods achieved comparable performance, the Shan–Bohn German equations consistently yielded lower or comparable errors for both males and females and provided a simpler implementation. We therefore adopted the Shan–Bohn German model as our literature-best baseline for subsequent comparisons and for partitioning lumped segment masses within our framework.

### Supplementary B. Geometric model

This section provides the analytical equations used to calculate segment volumes, centers of mass, and moments of inertia. All segment definitions are described in Section 3.1 of the main article. Shoulder depth, not reported in the ANSUR II dataset, was set equal to chest depth for BMI≤30 kg/m$^2$ and held constant above this threshold, as trunk enlargement with obesity primarily affects the lower torso. Neck circumference was included among the cross-sectional predictors to improve torso-segment estimations [7]. Mass redistribution during obese-specific adjustments excluded the hands and feet, which contain negligible adipose tissue [8]. Segment axes were aligned with anatomical directions (x: anteroposterior, y: vertical, z: mediolateral), and combined inertial properties were obtained using the parallel-axis theorem.

The segment volumes ($V$), centers of mass ($\mathbf{y}_{\mathrm{COM}}$), and principal moments of inertia ($I_{xx}, I_{yy}, I_{zz}$) were calculated analytically from the geometric model. Each body segment was represented by one or a combination of three geometric primitives (Fig. 2): (i) the head–neck and hand segments were modeled as ellipsoids; (ii) the lower torso and thigh segments included an elliptical cone; and (iii) the remaining parts were modeled as frustums of elliptical cones. The volume, COM position, and moments of inertia of an ellipsoid are calculated as

$$\begin{aligned}
&V = \tfrac{4}{3}\pi R^2 L,\\
&y_{\mathrm{COM}} = \tfrac{1}{2}L,\\
&I_{xx} = I_{zz} = \tfrac{1}{5}m(R^2 + L^2),\\
&I_{yy} = \tfrac{2}{5}mR^2.
\end{aligned}$$

where $R$ and $L$ denote the semi-axes along the radial and longitudinal directions, respectively, and $m$ is the segment mass. For an elliptical cone, the corresponding quantities are given by

$$\begin{aligned}
&V = \tfrac{1}{3}\pi LRr,\\
&y_{\mathrm{COM}} = \tfrac{3}{4}L,\\
&I_{xx} = \tfrac{3}{20}m(4R^2 + L^2),\\
&I_{yy} = \tfrac{3}{10}m(r^2 + R^2),\\
&I_{zz} = \tfrac{3}{20}m(4r^2 + L^2).
\end{aligned}$$

where $R$ and $r$ are the major and minor semi-axes at the base, and $L$ is the height of the cone. For a frustum of an elliptical cone, the equations (volume, COM, and moment of inertia at the upper cross) are

$$\begin{aligned}
&V = \tfrac{\pi L}{3}\left(R_1 r_1 + R_2 r_2 + \sqrt{R_1 r_1 R_2 r_2}\right),\\
&y_{\mathrm{COM}} = L\frac{r(R_1 + R_2) + r_2(R_1 + 3R_2)}{2r_1(2R_1 + R_2) + r_2(R_1 + 2R_2)}
\end{aligned}$$

$$\begin{aligned}
I_{xx} = \frac{\pi}{240}L\rho \left[4h^2\left(r_1(2R_1 + 3R_2) + 3r_2(R_1 + 4R_2)\right) + \right.\\
\left. 3\left(r_1\left(4R_1^2 + 3R_1^2R_2 + 2R_1R_2^2 + R_2^2\right) + r_2\left(R_1^2 + 2R_1^2R_2 + 3R_1R_2^2 + 4R_2^2\right)\right)\right],
\end{aligned} \tag{1}$$

$$\begin{aligned}
I_{yy} = \frac{\pi}{80}L\rho \left[r_1^4(4R_1 + R_2) + r_1^2 r_2(3R_1 + 2R_2) + \right.\\
r_1\left(4R_1^2 + 3R_1^2R_2 + 3r_2^2R_2 + R_2^2 + 2R_1(r_2^2 + R_2^2)\right) + \\
\left. r_2^2\left[R_1^2 + 2R_1^2R_2 + 4R_2(r_2^2 + R_2^2)\right] + R_1(r_2^2 + 3R_2^2)\right],
\end{aligned} \tag{2}$$

$$\begin{aligned}
I_{zz} = \frac{\pi}{240}L\rho \left[4h^2\left(r_1(2R_1 + 3R_2) + 3r_2(R_1 + 4R_2)\right) + \right.\\
\left. 3\left[r_1^4(4R_1 + R_2) + r_1^2 r_2(3R_1 + 2R_2) + r_1 r_2^2(2R_1 + 3R_2) + r_2^3(R_1 + 4R_2)\right]\right].
\end{aligned} \tag{3}$$

where $R_1, r_1$ and $R_2, r_2$ denote the major and minor semi-axes of the upper and lower cross-sections, respectively, $\rho$ indicates density, and $L$ is the longitudinal distance between them. All moments of inertia are expressed about the local segment coordinate frame aligned with anatomical directions (x: anteroposterior, y: vertical, z: mediolateral). For composite segments consisting of multiple primitives, combined inertial properties were obtained using the parallel-axis theorem.

### Supplementary C. Regression models

This section complements Sections 3.2–3.4 of the main manuscript and summarizes the regression models used to estimate segment lengths, cross-sectional parameters, and lumped segment masses. All models were trained using data from ANSUR II (for lengths and cross-sectional parameters) or NHANES (for segment masses), using height, weight, and sex as predictors. A range of regression approaches was evaluated, including linear models, regularized models (LAS and LASSO), support vector regression, and tree-based methods. The selected models for each parameter were chosen based on prediction accuracy and consistency across the BMI range.

Table S5 reports the selected regression model for each parameter and the associated prediction error (RMSE) for males and females. The table also indicates the input variables used for each regression.

**Table S5:** Selected regression models for segment length, mass, and cross-section parameters. The table presents the model with the lowest RMSE for each parameter. SVR: support vector regression with a linear kernel; LR: linear regression; LAS: LASSO; len: length; wd: width; dp: depth; circ: circumference.

| | **male** | **female** | | **male** | **female** | | **male** | **female** |
|---|---|---|---|---|---|---|---|---|
| **Length** (cm) *input = [height]* | | | **Circumference** (cm) *input = [height, weight]* | | | **Circumference** (cm) *input = [height, weight, hip circ, waist circ]* | | |
| head len | SVR 1.13 | LAS 1.03 | head circ | SVR 1.32 | SVR 1.44 | head circ | SVR 1.31 | SVR 1.44 |
| forearm len | LR 0.97 | SVR 0.89 | neck circ | LR 1.52 | LR 1.17 | neck circ | LR 1.41 | LR 1.10 |
| upperarm len | SVR 1.06 | SVR 0.99 | neck base circ | SVR 1.60 | LR 1.43 | neck base circ | SVR 1.57 | LR 1.38 |
| hand len | SVR 0.68 | SVR 0.68 | shoulder circ | SVR 3.23 | LR 2.73 | shoulder circ | SVR 3.09 | LR 2.64 |
| upper torso | LAS 1.76 | LR 2.03 | chest circ | LR 3.12 | SVR 4.60 | chest circ | LR 2.87 | SVR 4.01 |
| middle torso | LR 2.06 | LAS 2.18 | waist circ | LR 4.00 | LR 4.76 | biceps circ | LR 1.80 | LR 1.44 |
| lower torso | LR 2.29 | LR 2.11 | buttock circ | SVR 2.79 | LR 2.86 | forearm circ | LAS 1.24 | LR 1.14 |
| thigh len | SVR 2.00 | LAS 1.76 | biceps circ | LR 1.85 | LR 1.49 | hand circ | SVR 0.79 | SVR 0.70 |
| shank len | SVR 1.34 | LR 1.50 | forearm circ | LAS 1.24 | LR 1.14 | wrist circ | SVR 0.60 | SVR 0.51 |
| ankle hgt. | LR 0.47 | LR 0.43 | hand circ | SVR 0.79 | SVR 0.70 | thigh circ | SVR 2.30 | LR 2.18 |
| foot len | LR 0.86 | LR 0.80 | wrist circ | SVR 0.60 | SVR 0.51 | lower thigh circ | LR 1.54 | SVR 1.77 |
| biacromial wd | LR 1.54 | LR 1.38 | thigh circ | SVR 2.30 | LR 2.18 | heel-ankle circ | LR 0.97 | SVR 0.95 |
| head wd | LAS 0.64 | LAS 0.67 | lower thigh circ | LR 1.54 | SVR 1.77 | calf circ | LR 1.57 | LR 1.70 |
| shoulder-waist len | LR 2.33 | LAS 2.07 | heel-ankle circ | LR 0.97 | SVR 0.95 | ankle circ | LR 0.96 | SVR 1.08 |
| buttock hgt. | SVR 2.65 | SVR 2.36 | calf circ | LR 1.57 | LR 1.70 | ball of foot circ | LAS 1.01 | LR 0.91 |
| crotch hgt. | SVR 2.30 | SVR 2.23 | ankle circ | LR 0.96 | SVR 1.08 | chest wd | LR 1.22 | LR 1.26 |
| hand wd | LR 0.38 | SVR 0.32 | ball of foot circ | LAS 1.01 | LR 0.91 | hip wd | LR 1.17 | LR 1.36 |
| head wd | SVR 0.54 | SVR 0.48 | chest wd | LR 1.22 | LR 1.26 | waist wd | SVR 1.44 | LR 1.80 |
| **Mass** (kg) *input = [height, weight]* | | | waist dp | SVR 1.54 | SVR 1.57 | | | |
| leg mass | SVR 0.87 | LR 1.10 | chest dp | SVR 1.19 | SVR 1.56 | | | |
| arm mass | SVR 0.38 | SVR 0.33 | buttock dp | LR 1.11 | SVR 1.04 | | | |
| trunk mass | SVR 1.84 | SVR 2.01 | | | | | | |
| head mass | LAS 0.39 | LAS 0.34 | | | | | | |

### Supplementary D. Extended results

This section presents supplementary analyses conducted to validate key modeling assumptions in the framework. We evaluated whether additional anthropometric inputs improve segment-length predictions and examined how waist and hip circumferences capture principal modes of body-shape variation.

First, segment lengths were not meaningfully influenced by body weight, despite its frequent use in prior regression models [4, 6]. To verify this, we evaluated three progressively extended linear models (height only; height and weight; height, weight, and BMI) using identical three-fold GroupKFold cross-validation stratified by ethnicity. For each body segment, we computed the out-of-sample RMSE for all three models and quantified the paired difference $\Delta = \text{RMSE}_{\text{extended}} - \text{RMSE}_{\text{height-only}}$. Across all segments, the mean $\Delta$ values for both sexes were near zero, with 95% confidence intervals remaining within a $\pm 0.5$ mm non-inferiority margin. These results indicate that, once height is included as a predictor, neither body weight nor BMI provides meaningful additional predictive value for adult segment lengths.

Second, waist and hip circumferences emerged as key predictors of body-shape variation through the feature-selection analysis, consistent with their conventional use as descriptors of body-type categories (e.g., apple- and pear-shaped) [9]. We performed a supplementary Principal Component Analysis (PCA) of all standardized body circumferences in the ANSUR II dataset. The first principal component (PC1) explained the majority of inter-individual variance (67.1% in males; 61.9% in females) and reflected a global size/volume mode with uniformly positive loadings across girths. Predicting each principal component from waist and hip circumferences using five-fold GroupKFold cross-validation showed that PC1 was strongly explained ($R^2 \approx 0.8$), whereas higher-order components, each accounting for less than 10% additional variance, had near-zero predictive power ($R^2 \approx 0$).

[2] Centers for Disease Control and National Center for Health Statistics (NCHS) Prevention (CDC). National health and nutrition examination survey questionnaire. Hyattsville, MD: U.S. Department of Health and Human Services, Centers for Disease Control and Prevention.

[3] Renato Contini. Body segment parameters, part ii. *Artificial limbs*, 16(1):1–19, 1972.

[4] Gongbing Shan and Christiane Bohn. Anthropometrical data and coefficients of regression related to gender and race. *Applied ergonomics*, 34(4):327–337, 2003.

[5] Carol Ann Finch. Estimation of body segment parameters of college age females using a mathematical model. 1985.

[6] Vladimir Zatsiorsky. The mass and inertia characteristics of the main segments of the human body. *Biomechanics*, pages 1152–1159, 1983.

[7] Zachary Merrill, Subashan Perera, and Rakié Cham. Predictive regression modeling of body segment parameters using individual-based anthropometric measurements. *Journal of biomechanics*, 96:109349, 2019.

[8] K Kotani, K Tokunaga, S Fujioka, T Kobatake, Y Keno, S Yoshida, I Shimomura, S Tarui, and Y Matsuzawa. Sexual dimorphism of age-related changes in whole-body fat distribution in the obese. *International journal of obesity and related metabolic disorders: journal of the International Association for the Study of Obesity*, 18(4):207–202, 1994.

[9] Caroline M Apovian, Louis Aronne, and Sarah R Barenbaum. *Clinical management of obesity*. Professional Communications, Inc, 2023.
